# Transposable element dynamics are consistent across the *Drosophila* phylogeny, despite drastically differing content

**DOI:** 10.1101/651059

**Authors:** Tom Hill

## Abstract

**Background:** The evolutionary dynamics of transposable elements (TEs) vary across the tree of life and even between closely related species with similar ecologies. In *Drosophila*, most of the focus on TE dynamics has been completed in *Drosophila melanogaster* and the overall pattern indicates that TEs show an excess of low frequency insertions, consistent with their frequent turn over and high fitness cost in the genome. Outside of *D. melanogaster*, insertions in the species *Drosophila algonquin*, suggests that this situation may not be universal, even within *Drosophila*. Here we test whether the pattern observed in *D. melanogaster* is similar across five *Drosophila* species that share a common ancestor more than fifty million years ago.

**Results:** For the most part, TE family and order insertion frequency patterns are broadly conserved between species, supporting the idea that TEs have invaded species recently, are mostly costly and dynamics are conserved in orthologous regions of the host genome

**Conclusions:** Most TEs retain similar activities and fitness costs across the *Drosophila* phylogeny, suggesting little evidence of drift in the dynamics of TEs across the phylogeny, and that most TEs have invaded species recently.

## Introduction

Transposable elements are selfish mobile genetic elements found throughout the genomes of most living organisms; these sequences copy and move throughout hosts genomes, mostly to the detriment of the host [1–5]. Mammalian genomes are rarely invaded by TEs, and therefore have few active transposable elements (TEs), a large proportion of their genomes are composed of TE insertions fixed within a species population [6–8]. Comparatively, TEs in the fruit fly *Drosophila* appear to be highly active, resulting in polymorphic insertions for most TE families within a species population, with a lower proportion of their genome comprised of TEs [3, 9].

These differences can be explained with a model described by Lee and Langley [10]. TEs have bursts of activity recently after invading a genome, resulting in their transposition, their insertions are primarily deleterious to the host; they can interrupt a gene, cause aberrant expression or differential exon expression [3, 4, 9, 11]. Without regulation, TEs are also rampantly expressed and transposing, at a high cost to the host [10, 12]. To combat this, TE activity is suppressed, in the case of most animals, via the piRNA system [13–16]. Using small RNAs transcribed from TE sequences, the piRNA system targets and degrades complementary TE mRNAs and cause heterochromatin formation on similar TE insertions [12, 17–19]. However, the piRNA system can cause the propagation of heterochromatic silencing marks around TE insertions, resulting in the silencing of nearby genes and position effect variegation [10, 19]. This deleterious side effect, in combination with the deleterious effects of TE insertions suggests TE insertions should be rare in euchromatic regions [3, 9, 10, 20].

Within this model, TEs will enter a genome and spread rapidly through a burst of unsuppressed transposition [21–23]. The TE will be silenced via the piRNA system and regulated so long as piRNAs are produced against the TE [12, 24]. Following this, a TE will decrease in activity and will have an insertion frequency spectra (IFS) changing from an excess of rare insertions to fewer, more common insertions, likely in heterochromatic regions and piRNA clusters [11, 13], as the TE ages [22]. From this, we expect larger genomes with fewer active TEs, such as mammals, to have higher TE abundances and TE insertion frequency spectra showing no skew towards rare insertions as TE insertions are on average, less costly as genes make up a smaller portion of the genome, and less likely to have bursts of activity (Figure 1). While species with higher effective population sizes, higher coding densities and more active (more recently invaded) TEs, such as *Drosophila melanogaster*, should have lower abundances of TEs and IFS skewed to rare insertions [10, 11, 21, 22, 25].

**Figure 1:**
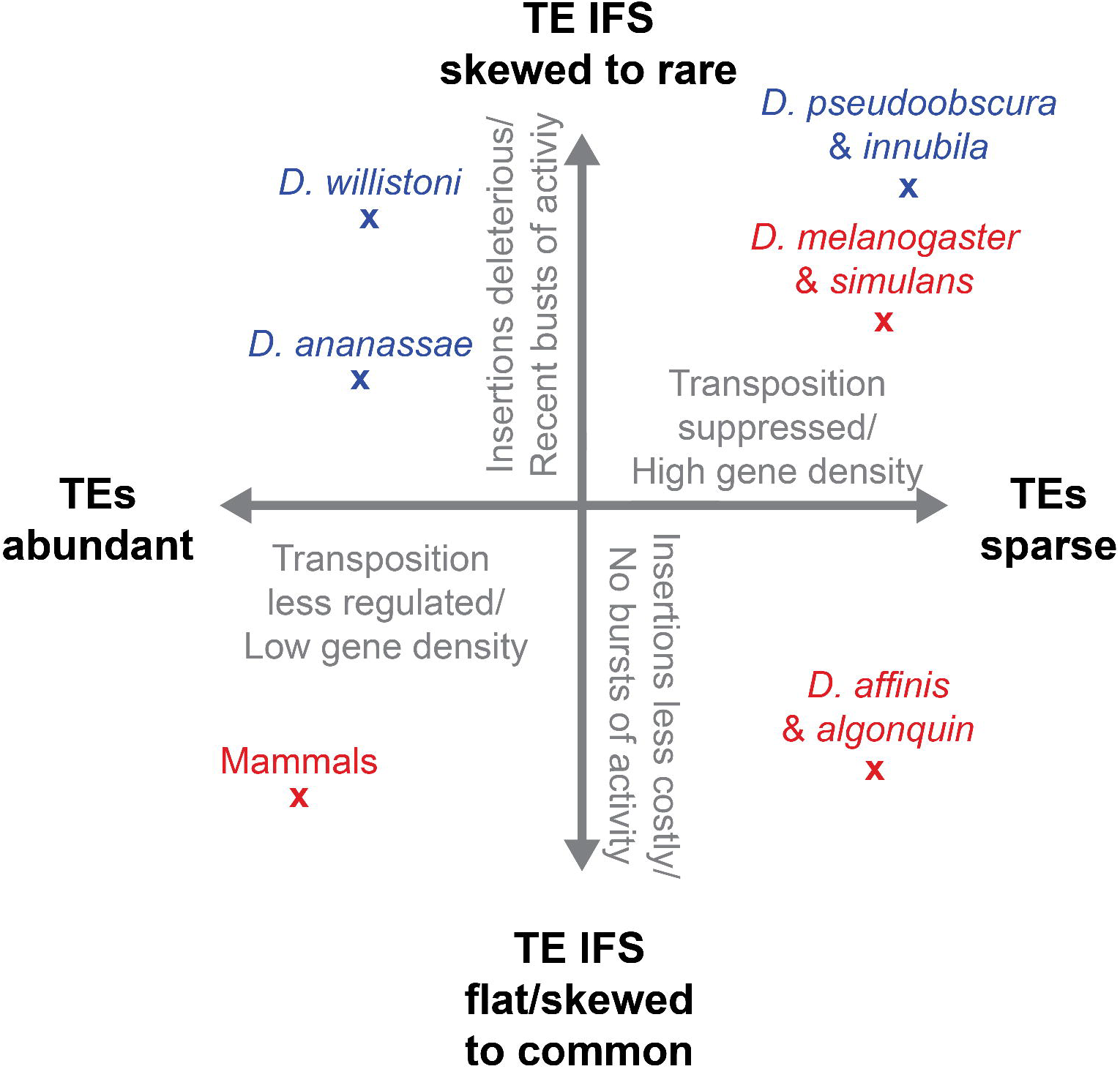
Schematic depicting the model explaining the differences in TE abundance and insertion frequency spectra across species, with species analyzed previously in red, species analyzed here in blue. Species have been placed in the schematic based on 1 – the insertion frequency spectrum relative to mammals and *D. melanogaster*, and 2 – TE abundances compared to mammals and *D. melanogaster*.

However, the expectation of lower euchromatic TE abundances, consistent with higher coding densities seen in *Drosophila melanogaster* is not seen in all *Drosophila* species [23]. The dynamic nature of *Drosophila* TEs can be clearly seen in the 12-genomes project, a group of 12 sequenced *Drosophila* species genomes, that span the ~50 million year *Drosophila* genus, with species in both the *Drosophila* and *Sophophora* sister subgenera [26, 27]. The sequenced species show striking differences between TE families and orders, and make up differing proportions of the genome, between 5 and 40% across the tree [28]. Furthermore, two studies comparing *D. melanogaster* and *D. simulans* TE content find a divergence in activity between genomes, specifically finding higher TE content in *D. melanogaster*, an relatively more intermediate frequency insertions in *D. simulans*, and an relatively more fixed and low frequency insertions in *D. melanogaster* [22, 29, 30]. These studies support the idea that even the same families shared between closely related species can diverge in activity and insertion frequency over time, and that differences may be even more extremely between more diverged species [22, 29]. Finally, the TE content of two species in the *D. affinis* subgroup, is not comprised of lower copy number families with an excess of low frequency insertions [31]. Instead they have a few, highly abundant families, with many high frequency insertions, like mammalian genomes, despite their small genome and large effective population sizes [32, 33]. This could be as these species lack any TEs that have recently invaded the genome and therefore have bursts of activity within the genome [22]. Though the methods used in this study are not truly comparable to modern techniques of assessing TE abundances, together with the diversity of abundances in the 12 genomes it brings into question the extent to which the previously described model fits outside the *D. melanogaster*, and where within the frame work other species fit [23, 31].

Here, we use next generation sequencing data and modern TE content identification methods to assess the TE insertion densities and TE insertion frequency spectra of the euchromatic genome of five *Drosophila* species. We attempt to identify if TEs show patterns consistent with highly active TE families across species, suggested by insertions being rare and primarily deleterious and differ in their ages within species. Additionally, we examine the extent that TE insertion frequency patterns differ between species with differing abundances of TEs. We find that despite differences in TE abundances and euchromatic insertion densities between species, most TE insertions have an IFS consistent with families recently invading genomes, highly active and deleterious in all species, though some individual families differ in their insertion frequencies between species (Figure 1 and 3). This suggests that TEs remain consistently deleterious across the *Drosophila* phylogeny, despite strong phylogenetic differences between species, and large changes in effective population size and TE densities [28].

## Results

### TE content differs drastically across the species examined

To examine the abundance and fitness cost of TE insertions across our *Drosophila* phylogeny of five species (Figure 1, 2A), we sequenced 20 wild caught *Drosophila innubila* individuals (described in the materials and methods) and downloaded short-read data for 17 *D. melanogaster* genomes, 45 *D. pseudoobscura* Muller C’s, 16 *D. annanassae* genomes and 14 *D. willistoni* genomes. We then generated profiles of the TE content of each individuals in each species using a combination of *RepeatMasker*, *BEDTools* and *PopoolationTE2* (Tarailo-Graovac and Chen 2009; Quinlan and Hall 2010; Kofler *et al.* 2011b) [34–38] using either a repeat library generated from RepBase TE sequences (Data S1), or a custom repeat library for *D. innubila*, generated by RepeatModeler [39, 40]. We estimated the proportion of each genome made up of TE insertions, the median copy number of each TE family and the median insertion number of each family in the euchromatic portion of the genome. We grouped families by their orders, either terminal inverted repeat (TIR) and rolling circle (RC) DNA transposons, or long terminal repeat (LTR) and long interspersed nuclear elements (LINE) RNA retrotransposons [5, 41] (TE hierarchy in Data S2). Within each species, the TE content varies drastically – between 15% and 40% of each genome (Figure 2B), with consistently different numbers of TE copies and euchromatic insertions between species (Figure 2B). As identified elsewhere, there is a significant association between genome size and TE content (Table S2, *p*-value = 0.002) [5, 8, 42].

**Figure 2.**
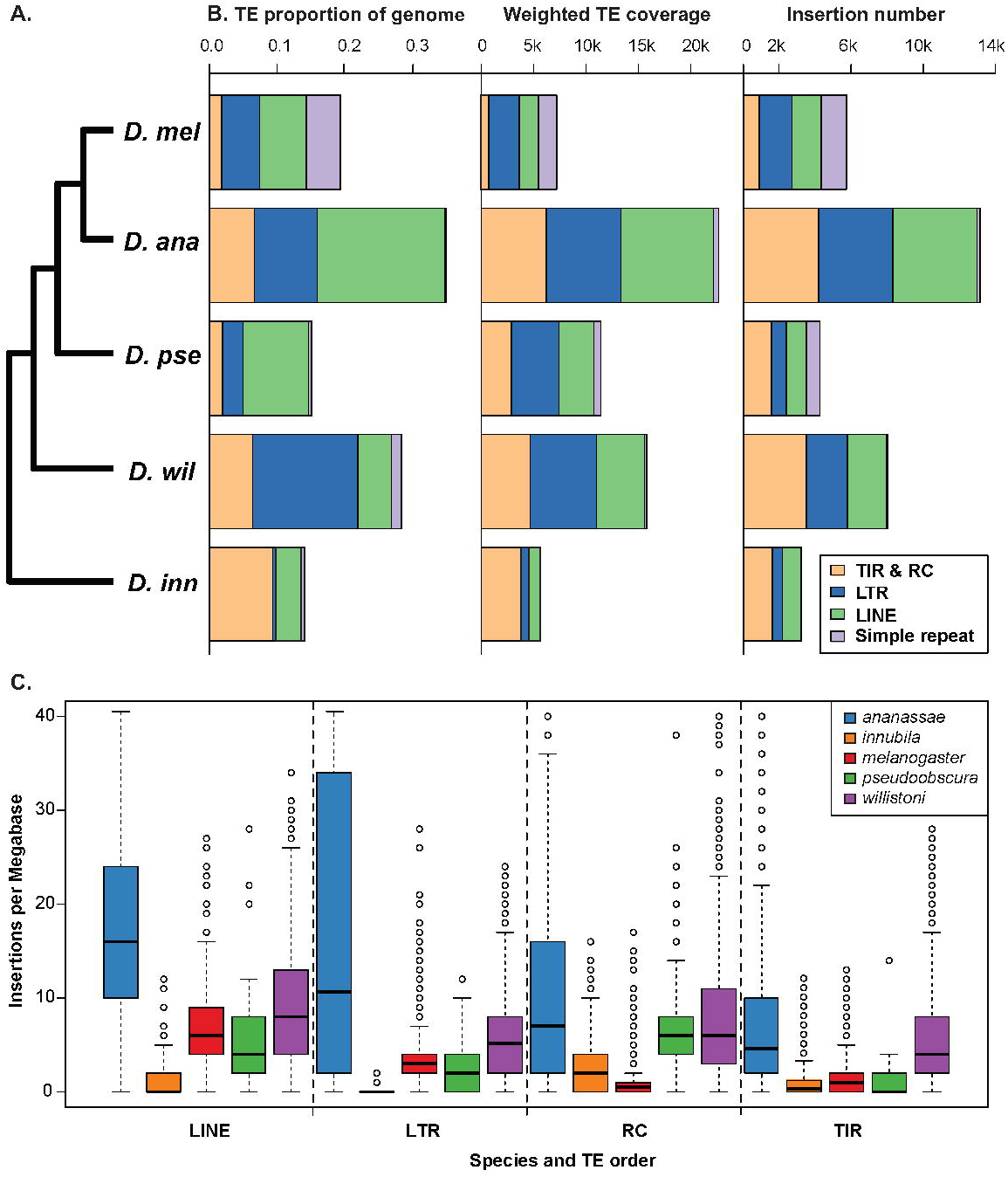
Transposable Element content (separated by TE order) in populations of five *Drosophila* species. TE content shown as **A.** Cartoon of tree of species assessed here, branches do not accurately represent the distance between species. **B.** Estimated TE profiles including TE proportions of each genome, median TE coverage, weighted by median nuclear coverage, and median TE insertion number. TIR and RCs were combined due to small numbers of either for many species. **C.** TE density per 1 Mb windows across the genome for each species and TE order.

The recently assembled and annotated genome of *D. innubila* has considerably lower insertion count numbers, perhaps due to the inferior annotation of TE content compared to other species. Interestingly, the *D. innubila* genome appears to have a lower amount of LTRs than most other studied *Drosophila* species [39], showing a similar profile to the relatively closely related *D. mojavensis* [28]. Most other species have retrotransposons, such as LTRs and LINEs, making up a large proportion of their repeat content (Figure 2B) [23]. As shown previously, *D. ananassae* and *D. willistoni* have much higher TE content than the other species analyzed here [23, 28]. These species differ in genome size, including an expanded Muller Element F in *D. ananassae* [23, 43]. In fact, there is an excess of TE content in *D. ananassae* on Muller element F. This Muller element represents only ~11.6% of the assembled reference genome (based on *D. melanogaster* orthology) but contains ~21.1% of the reference genomes TE content (based on *RepeatMasker* estimates), and so may account for the differences seen here.

To control for this Muller element expansion and other differences in genome size, we measured the TE insertion density per autosomal euchromatic megabase and found a significant excess of TE insertions per MB in *D. ananassae* and *D. willistoni* versus all other species, in all TE orders (Figure 2C, quasi-Poisson GLM, z-value > 19.296, *p*-value < 0.0006). These differences in TE abundances suggest that TE insertions may have differing dynamics between species, even when excluding TE rich regions. Due to the larger genomes and more abundant TE insertions, insertions may be less costly in *D. ananassae* and *D. willistoni* compared to other species and so may be more common in populations, with IFS skewed towards higher frequencies [12, 15, 44].

### TE insertions are primarily rare across the *Drosophila* phylogeny

Using the TE insertions called with *PopoolationTE2*, we found the insertion frequency spectrum (IFS) across each TE order, across all species, limited to the autosomes (Kofler *et al.* 2016). Like the differing TE insertion numbers and densities across species (Figure 2), the IFS also differ (Figure S1, Table S2 and S3). Comparing IFSs between TE orders, we find a significant excess of high frequency RC insertions in *D. melanogaster* versus other species (GLM quasi-Binomial *p*-value < 3.5e-5, t-value > 4.151). We also find an excess of rare (low frequency) TIR insertions versus other species in *D. innubila* (*p*-value = 2.37e-5, t-value = −4.24) and *D. pseudoobscura* (*p*-value = 5.74e-15, t-value = −7.891). Additionally, we find a significant excess of high frequency LTR insertions in *D. ananassae* versus all other species (GLM *p*-value < 2e-16, t-value = 13.243) and an excess of higher frequency LINE insertions in both *D. melanogaster* (GLM *p*-value < 2e-16, t = 12.526) and *D. ananassae* (GLM *p*-value < 2e-16, t-value =11.505). While we find IFS differ between species, in all cases TEs are skewed towards rare insertions (Figure 1). The median insertion frequency is below 25% in every TE order across all species and shows no significant differences between species (Table S2 and S3, GLM *p*-value > 0.213).

As these comparisons may be biased by factors such as how the data was generated, the sequencing methods, the quality of the reference genomes and the TE annotation, we limited our analysis to *D. melanogaster*, *D. ananassae* and *D. willistoni*, three species with data generated in similar manners, with similar TE families and high-quality reference genomes. We assessed only insertions in regions of the autosomal genome identified as orthologous using *progressiveMauve* [45]. When comparing the insertions in these orthologous regions, for all comparisons we find the TE dynamics are more consistent between species, with no significant differences in any comparison (Table S2, Figure 3, Figure S1B: GLM *p*-value > 0.21, t-value < 1.556).

**Figure 3:**
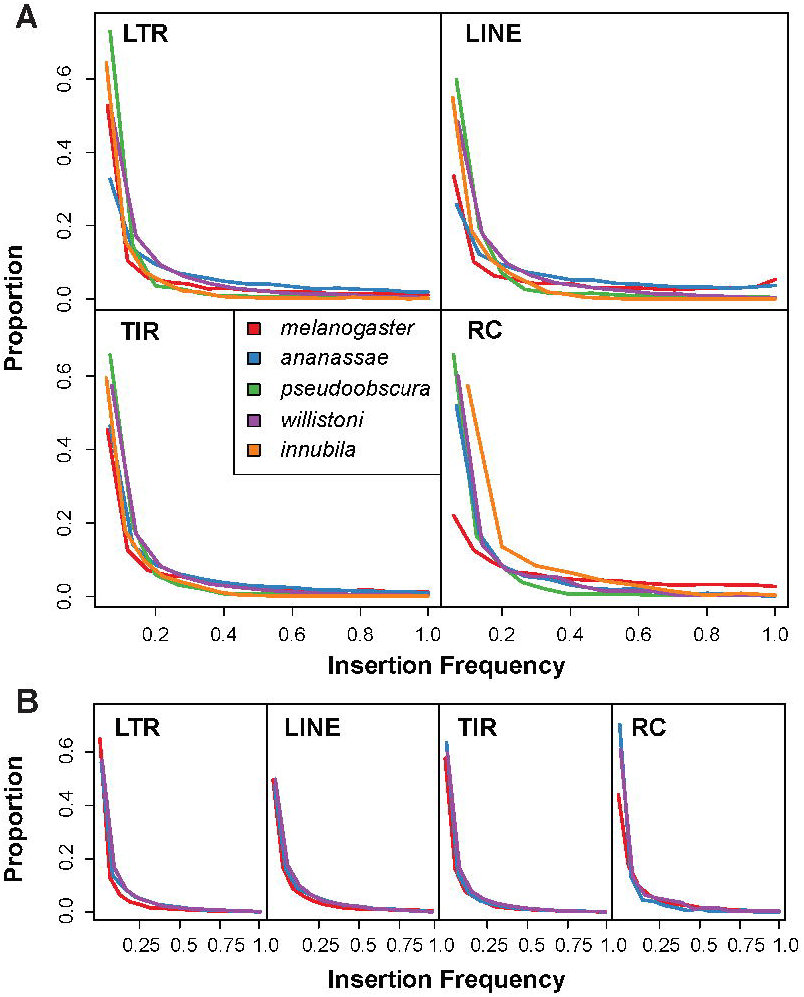
Average insertion frequency spectra for each species, separated by TE order for, A. TE insertions found across total genomes of all species. B. TE insertions called in orthologous regions for *D. melanogaster*, *D. willistoni* and *D. ananassae*.

### TE site frequency spectra rarely differ when accounting for population structure, insertions are primarily rare

One limitation of the analysis thus far is that all samples except *D. melanogaster* violate our implicit assumption of a single, panmictic population, which may skew the IFS to higher frequencies. This is can be seen in differences in estimated nucleotide site frequency spectrum of each species (limited to Muller element C for *D. pseudoobscura*) [46, 47], specifically finding an excess of high frequency variants in *D. pseudoobscura* when compared to *D. melanogaster* and an excess of low frequency variants in *D. willistoni* and *D. innubila* when compared to *D. melanogaster* (Figure S2, GLM quasi-Binomial *p*-value < 0.05). As expected, all site frequency spectra (SFS) show an excess of rare variants consistent with purifying selection, however *D. pseudoobscura* almost fits the neutral expectation, possibly due to the structured populations expected with the segregating inversions found on Muller element C [47–50].

To combat this, we clustered lines based on nuclear polymorphism using a principle component analysis (Figure S3). We then took a subset of lines for each species which appear to cluster as a single group in a principle component analysis (Figure S3). We also attempted to account for effective population size, on TE content, we find no association between effective population size and total TE content or insertion density, so did not control for this further (LM *p*-value > 0.05, Figure S3). This result contrasts with previous work which finds a negative association between effective population size and repetitive content [42, 51], possibly due to half of our species (*D. ananassae* and *D. willistoni*) as known exceptions for *Drosophila* TE content [23, 43]. Additionally, our sample size may be too small, and species too closely related to draw any strong conclusions about the relationship between repetitive content, genome size and effective population size in *Drosophila*.

In selected subpopulations, we examined the nuclear SFS between species and, with no drastic differences seen, we compared IFS between species. We find similar IFS across TE orders, though we do find an excess of high frequency RC insertions in *D. melanogaster* and an excess of high frequency LTR and LINE insertions in *D. ananassae* (Figure 3A, GLM *p*-value = 2e-16). As few RC insertions are found in *D. melanogaster* compared to other species, their genome has likely not been invaded by RC families recently, with their decreasing activity and increase in insertion frequency over time, while more recent invading RC families are found in other species are highly active with an excess of low frequency insertions [5, 22]. Again, we find no significant differences when comparing orthologous regions (GLM *p*-value > 0.05). As orthologous regions are exclusively in the euchromatic portion of the genomes, looking in orthologous regions removes heterochromatic insertions which are more likely to be high frequency [13]. As previous, most TE insertions are rare in all species (median frequency < 20%), with *D. ananassae* and *D. melanogaster* having the highest median frequency insertion, we also find no significant difference between median insertion frequency for any species or TE order (GLM *p*-value > 0.352) and no association between TE density or genome size with median insertion frequency (*p*-value > 0.05).

### Only a few, highly active, families differ across species, consistent with differing times of invasion

Our broader comparisons fit with previous work, that finds most TE families are highly active across a range of species due to recent spread of these TEs [22, 25]. As these broad observations may homogenize large differences between TE families, we chose to focus our analysis on specific families, shared between species.

We repeated the previous analysis across 10 TE super families found in all species [5, 23]. While there is a noticeable excess of low frequency insertions in *D. pseudoobscura*, we found no significant difference of insertion frequency between species for TE super family frequency (GLM logistic regression: −1.351 < *t*-value < −0.092, *p*-value > 0.183), however this may be due to few TE insertions in each subgroup or could again be too broad for any real inference (Figure S4).

Thus, we attempted to compare the dynamics of specific families shared between *D. melanogaster*, *D. ananassae* and *D. willistoni*. We identified 55 families shared between these species and extracted insertions for each family within the previously identified orthologous regions. For each TE family we compared the insertion frequency spectra for each species. Only eight of the 55 TE families showing any significant differences in IFS (six after multiple testing correction, Table S3-5, Figure S5, GLM logistic regression: *p*-value < 0.05). For these elements, one species has an excess of low frequency variants compared to the other two species (Figure S5), suggesting this difference may be due to a more recent acquisition in this species, resulting in higher activity, or more bursts of activity, rather than a consistent difference in activity between species, similar to the findings in comparisons across *D. melanogaster* group populations [22, 25, 52, 53].

To test if these elements more recently invaded the species showing a difference in dynamics, we calculated Tajima’s D for each of the shared 55 TE families. A negative Tajima’s D suggests an excess of low frequency variants, consistent with an expansion in copy number following a bottleneck, as would happen with a recent horizontal invasion [54, 55]. Among the 55 shared families, we find ten TE families have significant differences in estimations of Tajima’s D between species (GLM p-value < 0.05). Only one of the eight TE families with a differing IFS, has a significantly negative Tajima’s D, the *P-*element [30]. *P*-element has a significantly different insertion frequency spectrum between species (GLM logistic regression: *p*-value < 0.05), and significantly lower Tajima’s D (GLM p-value < 0.05), due to its recent horizontal transfer to *D. melanogaster* from *D. willistoni* [56–58]. Overall these results suggest few TE families differ between species in activity, after accounting for recent acquisitions.

## Discussion

Transposable elements, as mobile parasitic elements, are mostly costly to a host organism [3, 9, 20, 59], due to their rampant transposition, leading to the disruption of coding sequences [3, 9, 12, 20, 60], the mis regulation of gene expression [1, 2, 19, 61, 62] and even because of ectopic recombination and chromosomal breakage between two copies of the same TE family [3, 20, 63, 64]. Deleterious insertions are removed under purifying selection and TE families are rapidly silenced upon their acquisition [3, 11, 20, 64], giving an expectation for an insertion frequency spectrum skewed towards low frequency insertions for more recently acquired families that are highly active [3, 9, 20, 64, 65]. Most of the theoretical and experimental work that led to our understanding of TE dynamics has been completed in *D. melanogaster* [3, 20, 64, 66], under the assumption that TEs in other *Drosophila* and insects behave in a similar manner, despite some evidence to the contrary [31, 67, 68]. Here we test the validity of this assumption by assessing the TE dynamics in a *D. melanogaster* population and populations of four other *Drosophila* species. Despite drastic differences in TE content and densities between the species (Figure 2), we observe a pattern of rare insertions across all species, consistent with a recent invasion of highly active TE families, with insertions under strong purifying selection in all species (Figure 3, Figure S1, Table S2 and S4).

There are several possible explanations for the fact that work predating next generation sequencing technologies suggested differences in TE dynamics among species [31]. First, these differences may be due to host-specific factors (Table S2 −4, Figure S1 and S4), such as how recent the TE families have invaded a species genome (Hey 1989; Kaminker *et al.* 2002b). However, most families found in *D. affinis* are shared across the *D. obscura* group or with other *Drosophila* [69], suggesting TEs are being transferred between species. Second, high copy number families identified by *In Situ* hybridisation may have be low resolution conflating separate insertions as the same insertion, artificially inflating that insertion’s frequency and skewing its frequency higher than in lower copy number samples [31]. Finally, species genomes may differ in their chromatin states at different parts of genomes, limiting our analyses to well described euchromatic portions could stunt our ability to identify the diversity of TE dynamics in these host species. *D. ananassae*, for example, has an expansive Muller element F [43], full of transposable elements that was not included in this survey (due to most the chromosome being masked in the reference genome).

Overall, our results support a model where TE families invade genomes, expand in copy number, are rapidly regulated by the host genome (to differing levels among species), with insertions primarily being deleterious in all species examined, though the selection against insertions appears to differ from species to species to a minor degree.

## Materials and Methods

### Population genomic data

We used next generation sequencing data from five species collected from three sources, summarized in Table S1. For *Drosophila melanogaster*, we downloaded the FastQ files of 100bp paired end reads for a randomly selected set of 17 lines of the DPGP from a population collected from Zambia (SRA accessions: SRR203500-10, SRR204006-12). Similarly, we downloaded the FastQ files of 100bp paired end reads for 45 *Drosophila pseudoobscura* lines (SRA accessions: SRR617430-74) [47]. These lines consist of wild flies crossed to balancer stocks for chromosome 3 (Muller element C), this results in an isolated wild third chromosome, but a mosaic of balancer and wild stocks across the remainder of the genome, due to this we restricted our analysis to Muller element C (chromosome 3) in these lines.

We obtained sequencing information for 16 *Drosophila ananassae* isofemale lines and 14 *willistoni* isofemale lines. These lines were sequenced using an illumina HiSeq 2500 to produce 100bp paired end reads for each isofemale line.

Wild *Drosophila innubila* were captured at the Southwest Research Station in the Chiricahua Mountains between September 8^th^ and 15^th^, 2016. Baits consisted of store-bought white button mushrooms (*Agaricus bisporus*) placed in large piles about 30cm in diameter. A sweep net was used to collect the flies over the baits. Flies were sorted by sex and species at the University of Arizona and males were frozen at −80 degrees C before being shipped on dry ice to Lawrence, KS. All *D. innubila* males were homogenized in 50 microliters of viral buffer (a media meant to preserve viral particles, taken from [70]) and half of the homogenate was used to extract DNA using the Qiagen Gentra Puregene Tissue kit (#158689, Germantown, Maryland, USA). We constructed a genomic DNA library using a modified version of the Nextera DNA Library Prep kit (#FC-121-1031, Illumina, Inc., San Diego, CA, USA) meant to conserve reagents [71]. We sequenced 20 male samples as a library on two lanes of an Illumina HiSeq 2500 System Rapid-Run to generate paired-end 150 base-pair reads (available at NCBI accession numbers SRR6033015 [72]).

We trimmed all data using *Sickle* (minimum length = 50, minimum quality = 20) before mapping, and removed adapter sequences using *Scythe* [73, 74].

### Custom reference genomes

We downloaded the latest *Flybase* reference genome (Flybase.org, as of December 2018) for *D. melanogaster*, *D. ananassae*, *D. pseudoobscura* and *D. willistoni*, and used the *D. innubila* reference genome available on NCBI (NCBI accession: SKCT00000000) [39, 75, 76].

For the released genomes (*D. melanogaster*, *D. ananassae*, *D. pseudoobscura* and *D. willistoni*), we identified and masked each reference genome using *RepeatMasker* (parameters: - pa 4 –s –gff –gccalc –nolow –norna –no_is) [36, 37], using a custom repeat library, consisting of *Repbase* TE sequences previously identified in each of the species examined here [77].

For. *D. innubila*, we generated a repeat library for the reference genome using *RepeatModeler* (parameters: - engine NCBI) [40]. Then, after identifying each family order by NCBI universal *BLAST* [78], used this library as the custom TE library for repeat masking as described above. To validate these *RepeatModeler* consensus sequences for *D. innubila*, we mapped Illumina data to the TE library and kept only TE sequences with at least 1x the genomic coverage across 80% of the sequence (BWA MEM, default parameters [79, 80]).

For each species, we then generated a custom reference genome required for the use of *PopoolationTE2* [34]. For this we merged the masked reference genome, the custom TE library used for masking and the genome TE sequences, extracted using *BEDTools* [38]. Next, as described in the *PopoolationTE2* manual, we generated a hierarchy for each genome which assigned each TE sequence (all consensus sequences and reference sequences) to a TE family and TE order as described in [5, 77], either terminal inverted repeat (TIR) and rolling circle (RC) DNA transposons, or long terminal repeat (LTR) and long interspersed nuclear element (LINE) RNA retrotransposons.

### TE content and copy number differences between genomes

We quantified the amount of TE content for all species in three ways: a) proportion of the reference genome masked with *RepeatMasker*, b) median insertion count of each TE family across all lines in a species and c) median insertion count of each family using *PopoolationTE2*. For b), we found the median coverage for each TE family and the median coverage masked nuclear genome using *BEDTools* (genomeCoverageBed) [38], we divided the median TE coverage by the median nuclear coverage (subsampled to 15x coverage) to find the copy number of each family. Then we calculated the median adjusted TE coverage across all lines for each species. For c), we calculated the median TE insertion count for each family in each species, based on TE insertions called using *PopoolationTE2*. To control for differences in genome size across euchromatic regions, we also calculated the insertions per 1 Megabase windows (sliding 250kbp) for each TE order in each line for each species, only for contigs greater than 100kbp with less than 60% of the window masked by *RepeatMasker* [36, 37].

### Calling transposable element insertions across genomes

To identify the TE insertions throughout the genome in each line for each species, we followed the recommended *PopoolationTE2* pipeline for each species (*sourceforge.net/p/popoolation-te2/wiki/Walkthrough/*) [34]. Though *PopoolationTE2* is designed for use with population pools, we used an adjusted method to call germline insertions in individuals. We subsampled each line to 15x average nuclear coverage and followed the pipeline with appropriate cutoffs to exclude most somatic transpositions (map-qual = 15, min-count = 5, min-distance = −200, max-distance = 500). *PopoolationTE2* gives an estimated frequency of the insertion based on coverage of the TE breakpoint versus the genomic coverage, here we used this as a support score for each TE insertion [34]. We removed insertions found exclusively in one line with lower than 50% frequency in an individual line, we then merged all remaining insertion files for each species. We also removed all insertions in regions with more than 60% of the Megabase window masked by *RepeatMasker* [36, 37], we also limited our analysis to scaffolds associated with autosomes in all species.

We used *BEDTools* [38] to estimate the frequencies of each family’s insertions across each species, combining TE insertions of the same family within 100bp of each other. We used a binomial GLM in R [81] to assess differences in insertion frequencies between species for each TE order, considering a significant effect of species compared to *D. melanogaster* for a p-value < 0.05 for each set of TE order insertion frequencies. If all species have a significant effect in a consistent direction, we consider this to be a significant effect of *D. melanogaster* on insertion frequency. We also compared the median insertion frequency across species and TE orders and again fit a GLM to compare in R [81].

For a less bias comparison of insertion frequency spectra, we limited our analyses to genomes with data generated in similar fashions (*D. melanogaster*, *D. ananassae, D. willistoni*), and to orthologous euchromatic regions of the genome. For this we used *progressiveMauve* to identify orthologous regions of each genome [45], then converted these regions into a bedfile and excluded regions below 100kb, with over 60% of bases masked. We excluded *D. innubila* from this comparison due to its high sequence divergence from all other species and difficulty in finding similar TE families in other species, and *D. pseudoobscura* as it only its Muller element C represented natural variation. We then extracted insertions found in the orthologous regions using *BEDTools [38]* to compare insertion frequency spectra in orthologous regions.

### Polymorphism and summary statistics across the host genome and TE sequences

We called polymorphism across the host nuclear genome using *GATK HaplotypeCaller* [82] for each host and found the nuclear site frequency spectrum for each species using this data, which we confirmed using *ANGSD* (folded spectra, bootstraps = 100, reference sequence given, ancestral sequence not used) [83]. ANGSD was also used to perform a principle component analysis between samples in each species to look for population substructure [83].

### Estimating the effective population size of species

We used the previously generated folded site-frequency spectra from *ANGSD* in *StairwayPlot* for *D. melanogaster*, *D. innubila*, *D. ananassae* and *D. willistoni* (excluding *D. pseudoobscura* due to the method of the data generation) [83, 84]. For each estimated effective population size back in time, we found the harmonic mean of the effective size in the past 100,000 years and took that as the average size for the line. We then compared the TE copy number estimations to effective population size.

### TE families with dynamics differing between species

We next wanted to identify TE families shared between species to identify differences in activity between species. We aligned families of the same superfamily (defined in the *Repbase* TE database [77]) from each species using *MAFFT* and considered families within 95% identity to be the same family in different species [85]. We then checked these matching TE families manually to make sure family groupings were correct and removed an error grouping of *D. ananassae P*-Galileo element with *D. willistoni* and *melanogaster P*-element. All other groupings appeared to make sense and sequence showed high levels of similarity. We then compared the site frequency spectrum of these species using a logistic regression GLM. We also tested for differences in population genetic statistics to assess if differences are due to the recent acquisition of a family in a species. We calculated Watterson’s theta, pairwise diversity and Tajima’s D using *Popoolation* and estimated TE family copy number based on coverage [53], then compared these statistics across family and species using a generalized linear model, noting significant interactions between species and TE family.

## Abbreviations

TE: transposable element
TIR: terminal inverted repeat
LTR: long terminal repeat
LINE: long interspersed nuclear element
RC: rolling circle
GLM: generalized linear model
IFS: insertion frequency spectra

## Declarations

### Ethics approval and consent to participate

Not applicable

### Consent for publication

Not applicable

### Funding

This work was supported by a postdoctoral fellowship from the Max Kade foundation (Austria), a K-INBRE postdoctoral grant (NIH Grant P20 GM103418) to TH and NIH Grant R00 GM114714 to Rob Unckless, funding TH.

#### Competing Interests

The author declares that they have no competing interests.

### Authors’ contributions

TH performed bioinformatics analysis, statistical analysis, wrote, read and approved the manuscript.

### Data availability

*D. pseudooscura* data available on NCBI SRA: SRR617430-SRR617474. *D. melanogaster* data available on NCBI SRA: SRR203500-10, SRR204006-12. *D. anannasae, D. willistoni* and *D. innubila* data will be made available upon publication. *Drosophila* genomes can be downloaded from Flybase.org or NCBI. All data is available upon request before acceptance.

## Acknowledgements

We are extremely grateful for the advice provided by R. Unckless and J. Blumenstiel, for providing the sequencing information, advice on analysis and the production of the manuscript. We are also grateful for helpful discussion provided by J. R. Chapman, A. J. Betancourt, C. Schlotterer, R. Kofler, and B. Charlesworth. Thanks for S. W. Schaeffer for providing the *D. pseudoobscura* data used in this survey and advice concerning how the data should be used. Finally, we thank two anonymous reviewers and an anonymous editor for their suggestions and comments on the manuscript which has greatly improved the writing, science and interpretation of results.

---

**Table S1:** Table of *Drosophila* strains used in this study, including information on species, collection location and SRA number.

**Table S2:** Comparison of TE insertion frequencies between species and the fit of GLMs at different levels showing significant differences between species.

**Table S3:** TE insertions across the analysed scaffolds for each of the five species analysed here, with TE family, superfamily, order and TE insertion site occupancy.

**Table S4:** TEs showing significant differences in distributions between species and the median Tajima’s D for each species to see if a recent horizontal acquisition was the cause of this difference. NA is given if the TE family is absent from the species in question.

**Table S5:** Table of GLM results for differences in IFS between TE families shared across *D. ananassae*, *melanogaster* and *willistoni* in shared regions of the genome.

**Figure S1. A.** Insertion frequency spectrum, plots showing the densities of insertions and the proportion of the population these insertions are found in. These spectra are estimated using *PopoolationTE2* for each species, separated by TE order. **B.** Insertion frequency spectrum of TE insertions for regions with high similarity, identified using *progressiveMauve*.

**Figure S2:** Site frequency spectra the nuclear genome of species analyzed here, calculated using ANGSD. The theoretical neutral site frequency spectrum is layered on top in red.

**Figure S3:** Principle component analysis for nuclear polymorphism for each species. Subpopulations are colored differently when known. E.G. Muller C inversion karyotype for *D. pseudoobscura* and Arizona sky island place of collection for *D. innubila* (both colored arbitrarily). Circled clusters are the lines used in the subset analysis, chosen arbitrarily based on the clustering seen in the PCAs. TE copy number for each species (+-2 * standard deviations) is also compared to estimated effective population size from *StairwayPlot*.

**Figure S4:** Insertion frequency per species for shared TE superfamilies’.

**Figure S5:** Site frequency spectrum of TEs shared between species that are significantly different in at least one comparison. Spectra are weighted by copy number. These are the 9 of 55 comparisons to show significant differences in distribution between species. The peak at ~60% in Harbinger-1 in *D. willistoni* is caused by a small number of insertions at 60% frequency and low insertion numbers found in the *D. willistoni*.

**Data S1:** TE sequences used in this study to determine repetitive content of each species.

**Data S2:** TE hierarchy for each species, separated by species. Hierarchy gives the specific sequence, the TE family it belongs to and its order.

## Notes

#### Summary of Updates

Minor revisions included from Mobile DNA review

## Bibliography

1. McClintock B: Induction of instability at selected loci in Maize. Genetics 1953, 38:579–599.

2. McClintock B: The origin and behavior of mutable loci in Maize. Proc Natl Acad of Sci USA 1950, 36:344–355.

3. Charlesworth B, Langley CH: The population genetics of *Drosophila* transposable elements. Annual review of genetics 1989, 23:251–287.

4. Burt A, Trivers R: Genes in Conflict. 2006.

5. Wicker T, Sabot F, Hua-Van A, Bennetzen JL, Capy P, Chalhoub B, Flavell A, Leroy P, Morgante M, Panaud O et al: A unified classification system for eukaryotic transposable elements. Nature reviews Genetics 2007, 8:973–982.

6. Hellen EHB, Brookfield JFY: The diversity of class II transposable elements in mammalian genomes has arisen from ancestral phylogenetic splits during ancient waves of proliferation through the genome. Molecular Biology and Evolution 2013, 30:100–108.

7. Hellen EHB, Brookfield JFY: Transposable element invasions. Mobile genetic elements 2013, 3:e23920.

8. Gregory TR: Synergy between sequence and size in large-scale genomics. Nature reviews Genetics 2005, 6:699–708.

9. Charlesworth B, Langley CH, Sniegowski PD: Transposable element distributions in *Drosophila*. Genetics 1997, 147:1993–1995.

10. Lee YCG, Langley CH: Transposable elements in natural populations of *Drosophila melanogaster*. Philosophical Transactions of the Royal Society B: Biological Sciences 2010, 365:1219–1228.

11. Lee YCG, Langley CH: Long-term and short-term evolutionary impacts of transposable elements on *Drosophila*. Genetics 2012, 192:1411–1432.

12. Blumenstiel JP: Evolutionary dynamics of transposable elements in a small RNA world. Trends in Genetics 2011, 27:23–31.

13. Lu J, Clark AG: Population dynamics of PIWI-interacting RNAs (piRNAs) and their targets in *Drosophila*. Genome Research 2010, 20:212–227.

14. Brennecke J, Aravin AA, Stark A, Dus M, Kellis M, Sachidanandam R, Hannon GJ: Discrete small RNA-generating loci as master regulators of transposon activity in *Drosophila*. Cell 2007, 128:1089–1103.

15. Aravin AA, Hannon GJ, Brennecke J: The piwi-piRNA pathway provides an adaptive defense in the transposon arms race. Science 2007, 318:761–764.

16. Brennecke J, Malone CD, Aravin AA, Sachidanandam R, Stark A, Hannon GJ: An epigenetic role for maternally inherited piRNAs in transposon silencing. Science (New York, NY) 2008, 322:1387–1392.

17. Senti KA, Jurczak D, Sachidanandam R, Brennecke J: piRNA-guided slicing of transposon transcripts enforces their transcriptional silencing via specifying the nuclear piRNA repertoire. Genes and Development 2015, 29:1747–1762.

18. Obbard DJ, Gordon KHJ, Buck AH, Jiggins FM: The evolution of RNAi as a defence against viruses and transposable elements. Philosophical transactions of the Royal Society of London Series B, Biological sciences 2009, 364:99–115.

19. Lee YCG: The role of piRNA-mediated epigenetic silencing in the population dynamics of transposable elements in *Drosophila melanogaster*. PLOS Genetics 2015, 11:1–24.

20. Langley CH, Montgomery E, Hudson R, Kaplan N, Charlesworth B: On the role of unequal exchange in the containment of transposable element copy number. Genetical Research 1988, 52:223–235.

21. Kofler R, Betancourt AJ, Schlötterer C: Sequencing of pooled DNA Samples (Pool-Seq) uncovers complex dynamics of transposable element insertions in *Drosophila melanogaster*. PloS Genetics 2012, 8:1–16.

22. Kofler R, Nolte V, Schlötterer C: Tempo and mode of transposable element activity in *Drosophila*. PLoS Genet 2015, 11:e1005406.

23. Clark AG, Eisen MB, Smith DR, Bergman CM, Oliver B, Markow Ta, Kaufman TC, Kellis M, Gelbart W, Iyer VN et al: Evolution of genes and genomes on the *Drosophila* phylogeny. Nature 2007, 450:203–218.

24. Senti KA, Brennecke J: The piRNA pathway: A fly’s perspective on the guardian of the genome. Trends in Genetics 2010, 26:499–509.

25. Petrov Da, Fiston-Lavier A-S, Lipatov M, Lenkov K, González J: Population genomics of transposable elements in *Drosophila melanogaster*. Molecular Biology and Evolution 2011, 28:1633–1644.

26. Markow TA, O’Grady P: *Drosophila*: a guide to species identification. 2006.

27. Markow TA, O’Grady PM: *Drosophila* biology in the genomic age. Genetics 2007, 177:1269–1276.

28. Sessegolo C, Burlet N, Haudry A, Biémont C, Vieira C, Tenaillon M, Hollister J, Gaut B, McClintock B, Mackay T et al: Strong phylogenetic inertia on genome size and transposable element content among 26 species of flies. Biology letters 2016, 12:521–524.

29. Adrion JR, Begun DJ, Hahn MW: Patterns of transposable element variation and clinality in *Drosophila*. Mol Ecol 2019, 28(6):1523–1536.

30. Kofler R, Hill T, Nolte V, Betancourt AJ, Schlötterer C: The recent invasion of natural *Drosophila simulans* populations by the P-element. Proceedings of the National Academy of Sciences 2015, 112:6659–6663.

31. Hey J: The transposable portion of the genome of *Drosophila algonquin* is very different from that in *Drosophila melanogaster*. Molecular Biology and Evolution 1989, 6:66–79.

32. Palmieri N, Kosiol C, Schlötterer C: The life cycle of *Drosophila* orphan genes. eLife 2014, 3:1–21.

33. McGaugh SE, Heil CSS, Manzano-Winkler B, Loewe L, Goldstein S, Himmel TL, Noor MaF: Recombination modulates how selection affects linked sites in *Drosophila*. PLoS biology 2012, 10:1–17.

34. Kofler R, Daniel G, Schl C: PoPoolationTE2: comparative population genomics of transposable elements using Pool-Seq. 2016:1–7.

35. Quesneville H, Bergman CM, Andrieu O, Autard D, Nouaud D, Ashburner M, Anxolabéhère D: Combined evidence annotation of transposable elements in genome sequences. PLoS Computational Biology 2005, 1:0166–0175.

36. Smit AFA, Hubley R: RepeatMasker Open-4.0; 2015.

37. Tarailo-Graovac M, Chen N: Using RepeatMasker to identify repetitive elements in genomic sequences. Current Protocols in Bioinformatics 2009.

38. Quinlan AR, Hall IM: BEDTools: a flexible suite of utilities for comparing genomic features. Bioinformatics (Oxford, England) 2010, 26:841–842.

39. Hill T, Koseva B, Unckless RL: The genome of *Drosophila innubila* reveals lineage-specific patterns of selection in immune genes. Molecular Biology and Evolution 2019:1–36.

40. Smit AFA, Hubley R: RepeatModeler Open-1.0. 2008.

41. Bao W, Kojima KK, Kohany O: Repbase Update, a database of repetitive elements in eukaryotic genomes. Mobile DNA 2015, 6:4–9.

42. Gregory TR, Johnston JS: Genome size diversity in the family *Drosophilidae*. Heredity 2008, 101:228–238.

43. Leung W, Students P: Retrotransposons Are the Major Contributors to the Expansion of the *Drosophila ananassae* Muller F. G3 2017, 7:2439–2460.

44. Levine MT, Malik HS: Learning to protect your genome on the fly. Cell 2011, 147:1440–1441.

45. Darling ACE, Mau B, Blattner FR, Perna NT: Mauve: Multiple Alignment of Conserved Genomic Sequence With Rearrangements. 2004:1394–1403.

46. Wallace AG, Detweiler D, Schaeffer SW: Evolutionary history of the third chromosome gene arrangements of *Drosophila pseudoobscura* inferred from inversion breakpoints. Molecular biology and evolution 2011, 28:2219–2229.

47. Fuller ZL, Haynes GD, Richards S, Schaeffer SW: Genomics of Natural Populations: How Differentially Expressed Genes Shape the Evolution of Chromosomal Inversions in. Genetics 2016.

48. Dobzhansky T: Genetics and the origin of species. 1937:364.

49. Dobzhansky T, Epling C: The suppression of crossing over in inversion heterozygotes of *Drosophila pseudoobscura*. Proceedings of the National Academy of Sciences of the United States of America 1948, 34:137–141.

50. Dobzhansky T, Sturtevant AH: Inversions In Chromosomes of *Drosophila pseudoobscura*. Genetics 1937, 23:28–64.

51. Lynch M, Conery JS: The Origins of Genome Complexity. Science 2003, 302:1401–1404.

52. Bergman CM, Haddrill PR: Strain-specific and pooled genome sequences for populations of *Drosophila melanogaster* from three continents. F1000 Research 2015, 31:1–5.

53. Kofler R, Orozco-terWengel P, de Maio N, Pandey RV, Nolte V, Futschik A, Kosiol C, Schlötterer C: Popoolation: A toolbox for population genetic analysis of next generation sequencing data from pooled individuals. PLoS ONE 2011, 6.

54. Tajima F: Statistical method for testing the neutral mutation hypothesis by DNA polymorphism. Genetics 1989, 123:585–595.

55. Bartolomé C, Bello X, Maside X: Widespread evidence for horizontal transfer of transposable elements across *Drosophila* genomes. Genome biology 2009, 10:R22.

56. Daniels SB, Peterson KR, Strausbaugh LD, Kidwell MG, Chovnick A: Evidence for horizontal transmission of the P transposable element between *Drosophila* species. Genetics 1990, 124:339–355.

57. Daniels SB, Strausbaugh LD: The distribution of P-element sequences in Drosophila: The willistoni and saltans species groups. 1986, 23:138–148.

58. Khurana JS, Wang J, Xu J, Koppetsch BS, Thomson TC, Nowosielska A, Li C, Zamore PD, Weng Z, Theurkauf WE: Adaptation to P-element transposon invasion in *Drosophila melanogaster*. Cell 2011, 147:1551–1563.

59. Doolittle WF, Sapienza C: Selfish genes, the phenotype paradigm and genome evolution. Nature 1980, 284:601–603.

60. Bachmann A, Knust E: The use of P-element transposons to generate transgenic flies. Methods in Molecular Biology 2008, 420:61–77.

61. Orgel LE, Crick FHC: Selfish DNA: the ultimate parasite. Nature 1980, 284:604–607.

62. Lisch D, Bennetzen JL: Transposable element origins of epigenetic gene regulation. Current Opinion in Plant Biology 2011, 14:156–161.

63. Sniegowski PD, Charlesworth B: Transposable element numbers in cosmopolitan inversions from a natural population of *Drosophila melanogaster*. Genetics 1994, 137:815–827.

64. Montgomery EA, Huang S, Langley CH, Judd BH: Chromosome rearrangement by ectopic recombination in *Drosophila melanogaster*: genome structure and evolution. Genetics 1991, 129:1085–1098.

65. Pasyukova EG, Nuzhdin SV, Morozova TV, Mackay TFC: Accumulation of transposable elements in the genome of *Drosophila melanogaster* is associated with a decrease in fitness. The Journal of heredity 2004, 95:284–290.

66. Petrov DA, Aminetzach YT, Davis JC, Bensasson D, Hirsh AE: Size matters: Non-LTR retrotransposable elements and ectopic recombination in *Drosophila*. Molecular Biology and Evolution 2003, 20:880–892.

67. Kaminker JS, Bergman CM, Kronmiller B, Svirskas R, Patel S, Frise E, David A, Lewis SE, Rubin GM, Ashburner M et al: The transposable elements of the *Drosophila melanogaster* euchromatin: a genomics perspective. Genome Biology 2002, 3:1–20.

68. Bergman CM, Bensasson D: Recent LTR retrotransposon insertion contrasts with waves of non-LTR insertion since speciation in *Drosophila melanogaster*. Proceedings of the National Academy of Sciences of the United States of America 2007, 104:11340–11345.

69. Hill T, Betancourt A: Extensive horizontal exchange of transposable elements in the *Drosophila pseudoobscura* group. Mobile DNA 2018, 20:1–14.

70. Nanda S, Jayan G, Voulgaropoulou F, Sierra-Honigmann AM, Uhlenhaut C, McWatters BJP, Patel A, Krause PR: Universal virus detection by degenerate-oligonucleotide primed polymerase chain reaction of purified viral nucleic acids. Journal of Virological Methods 2008, 152:18–24.

71. Baym M, Kryazhimskiy S, Lieberman TD, Chung H, Desai MM, Kishony R: Inexpensive multiplexed library preparation for megabase-sized genomes. PLoS ONE 2015:1–15.

72. Hill T, Unckless RL: The dynamic evolution of Drosophila innubila Nudivirus. Infection, Genetics and Evolution 2017:1–25.

73. Joshi N, Fass J: Sickle: A sliding window, adaptive, quality-based trimming tool for fastQ files. 2011, 1.33.

74. Buffalo V: Scythe; 2018.

75. Dos Santos G, Schroeder AJ, Goodman JL, Strelets VB, Crosby MA, Thurmond J, Emmert DB, Gelbart WM, Brown NH, Kaufman T et al: FlyBase: Introduction of the *Drosophila melanogaster* Release 6 reference genome assembly and large-scale migration of genome annotations. Nucleic Acids Research 2015, 43:D690–D697.

76. Gramates LS, Marygold SJ, Dos Santos G, Urbano JM, Antonazzo G, Matthews BB, Rey AJ, Tabone CJ, Crosby MA, Emmert DB et al: FlyBase at 25: Looking to the future. Nucleic Acids Research 2017, 45:D663–D671.

77. Kohany O, Gentles AJ, Hankus L, Jurka J: Annotation, submission and screening of repetitive elements in Repbase: RepbaseSubmitter and Censor. BMC bioinformatics 2006, 7:474.

78. Altschul SF, Gish W, Miller W, Myers EW, Lipman DJ: Basic local alignment search tool. Journal of Molecular Biology 1990, 215:403–410.

79. Li H, Handsaker B, Wysoker A, Fennell T, Ruan J, Homer N, Marth G, Abecasis GR, Durbin R: The sequence alignment/map format and SAMtools. Bioinformatics (Oxford, England) 2009, 25:2078–2079.

80. Li H, Durbin R: Fast and accurate short read alignment with Burrows-Wheeler transform. Bioinformatics (Oxford, England) 2009, 25:1754–1760.

81. Team RC: R: A Language and Environment for Statistical Computing. Vienna, Austria: R Foundation for Statistical Computing; 2013.

82. DePristo MA, Banks E, Poplin R, Garimella KV, Maguire JR, Hartl C, Philippakis AA, del Angel G, Rivas MA, Hanna M et al: A framework for variation discovery and genotyping using next-generation DNA sequencing data. Nature genetics 2011, 43:491–498.

83. Korneliussen TS, Albrechtsen A, Nielsen R: ANGSD: Analysis of Next Generation Sequencing Data. BMC Bioinformatics 2014, 15:356.

84. Liu X, Fu Y-X: Exploring population size changes using SNP frequency spectra. Nature genetics 2015, 47:555–559.

85. Katoh K, Misawa K, Kuma K-i, Miyata T: MAFFT: a novel method for rapid multiple sequence alignment based on fast Fourier transform. Nucleic acids research 2002, 30:3059–3066.

