## Supplementary Figures for "Transposable element dynamics are consistent across the *Drosophila* phylogeny, despite drastically differing content"

**Table S1:** Table of *Drosophila* strains used in this study, including information on species, collection location and SRA number.

**Table S2:** Comparison of TE insertion frequencies between species and the fit of GLMs at different levels showing significant differences between species.

**Table S3:** TE insertions across the analysed scaffolds for each of the five species analysed here, with TE family, superfamily, order and TE insertion site occupancy.

**Table S4:** TEs showing significant differences in distributions between species and the median Tajima’s D for each species to see if a recent horizontal acquisition was the cause of this difference. NA is given if the TE family is absent from the species in question.

**Table S5:** Table of GLM results for differences in IFS between TE families shared across *D. ananassae*, *melanogaster* and *willistoni* in shared regions of the genome.

**Figure S1. A.** Insertion frequency spectrum, plots showing the densities of insertions and the proportion of the population these insertions are found in. These spectra are estimated using *PopoolationTE2* for each species*,* separated by TE order. **B.** Insertion frequency spectrum of TE insertions for regions with high similarity, identified using *progressiveMauve*.

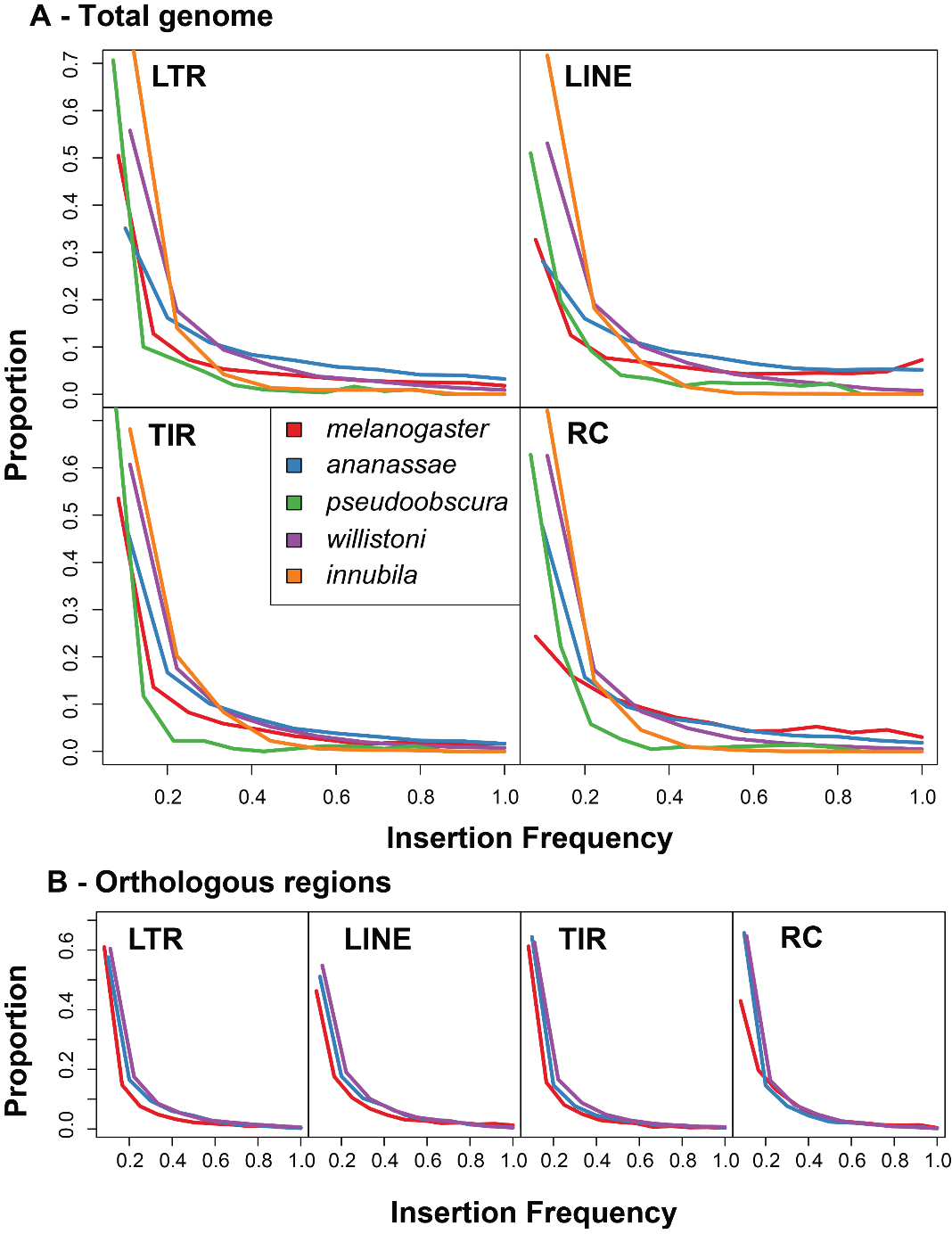

**Figure S2:** Site frequency spectra the nuclear genome of species analyzed here, calculated using ANGSD. The theoretical neutral site frequency spectrum is layered on top in red.

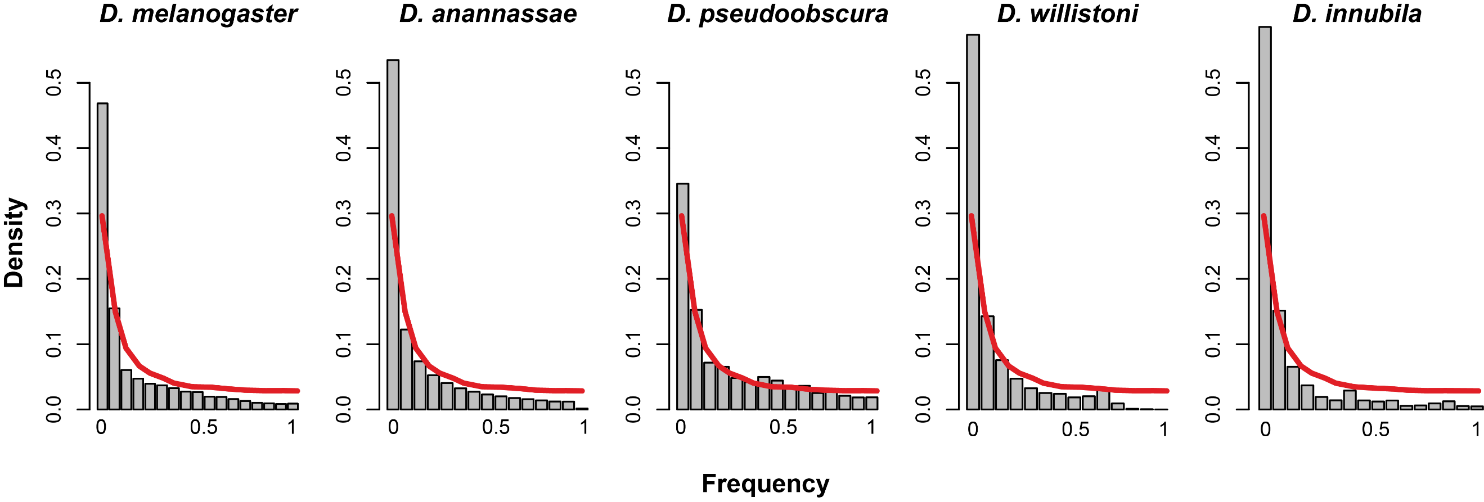

**Figure S3:** Principle component analysis for nuclear polymorphism for each species. Subpopulations are colored differently when known. E.G. Muller C inversion karyotype for *D. pseudoobscura* and Arizona sky island place of collection for *D. innubila* (both colored arbitrarily). Circled clusters are the lines used in the subset analysis, chosen arbitrarily based on the clustering seen in the PCAs. TE copy number for each species (+- 2 * standard deviations) is also compared to the harmonic mean of the estimated effective population size from *StairwayPlot*.

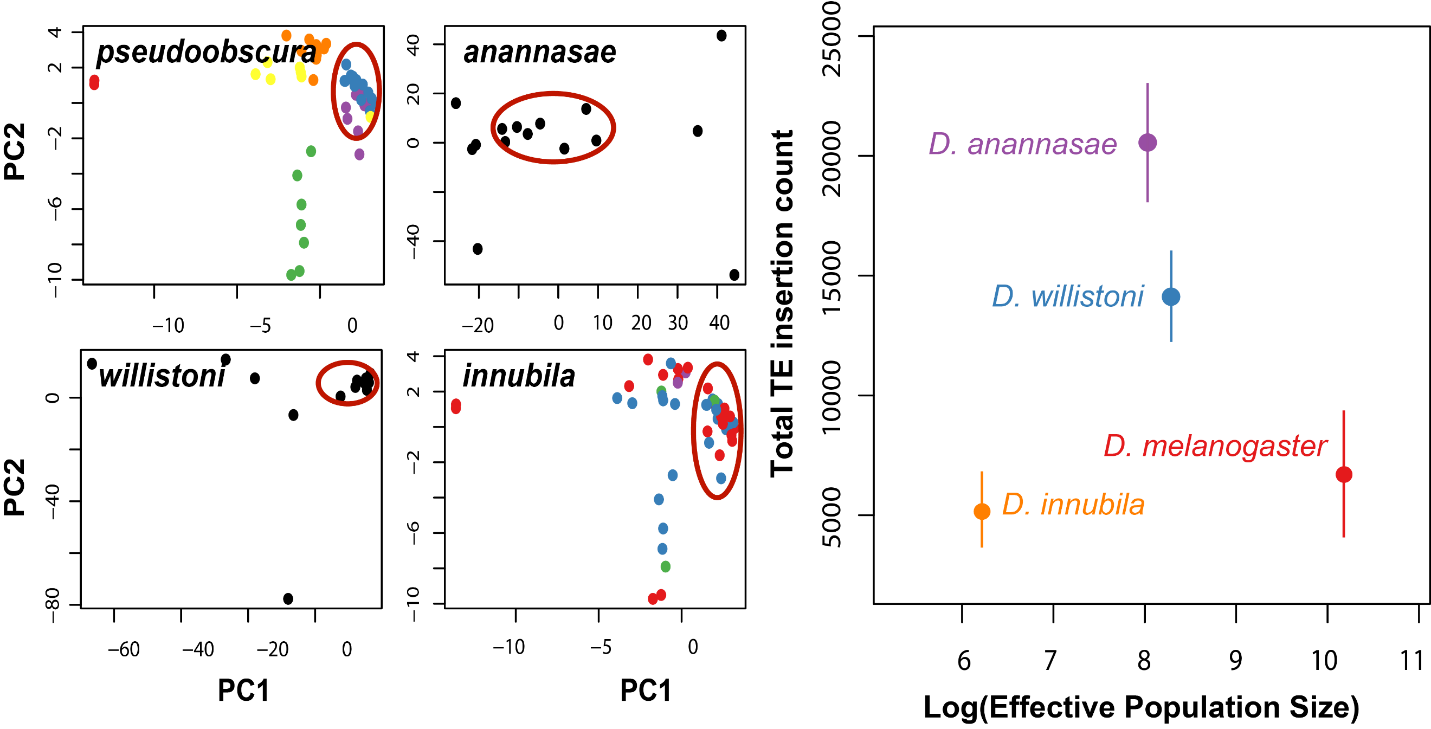

**Figure S4:** Insertion frequency per species for shared TE superfamily’s.

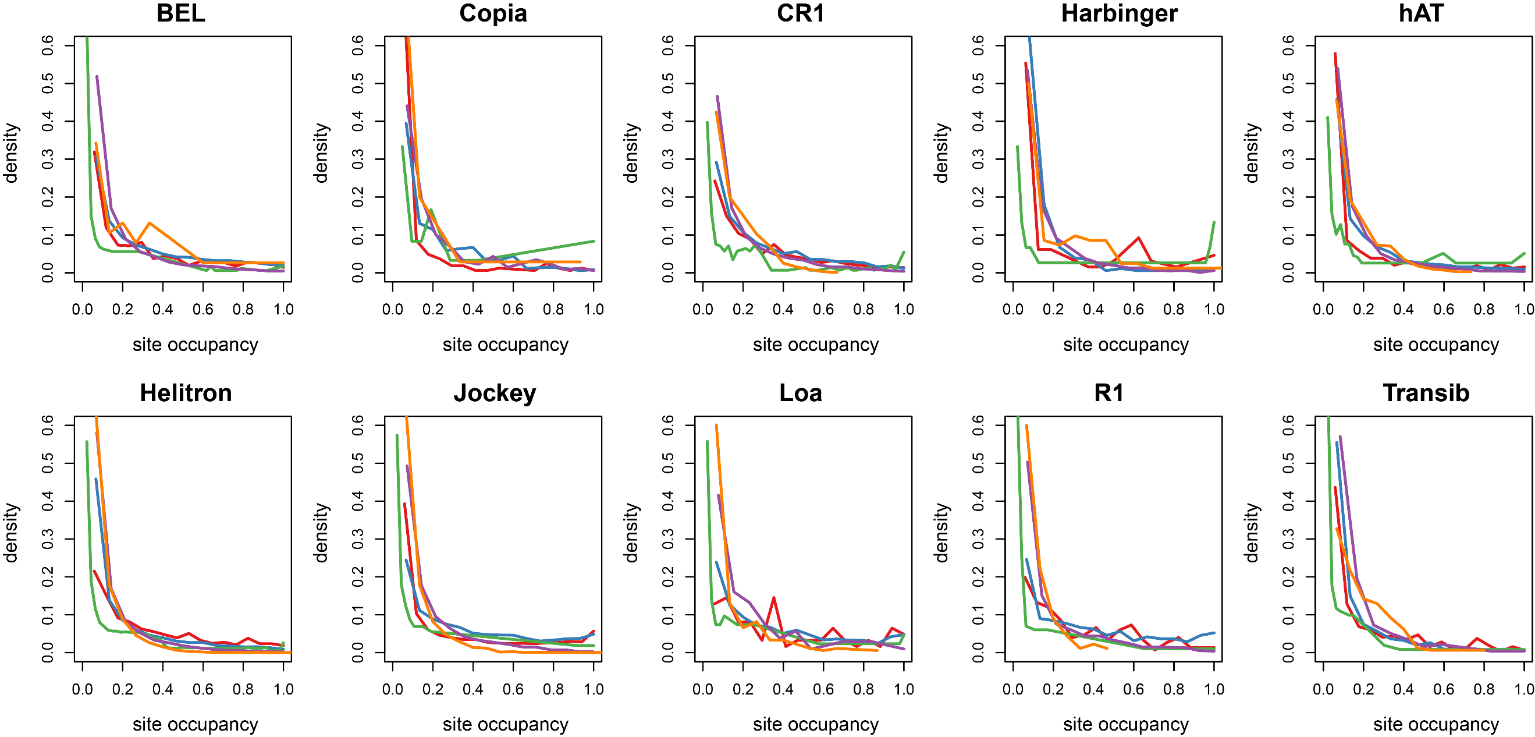

**Figure S5:** Site frequency spectrum of TEs shared between species that are significantly different in at least one comparison. Spectra are weighted by copy number. These are the 9 of 55 comparisons to show significant differences in distribution between species. The peak at ~60% in Harbinger-1 in *D. willistoni* is caused by a small number of insertions at 60% frequency and low insertion numbers found in the *D. willistoni*.

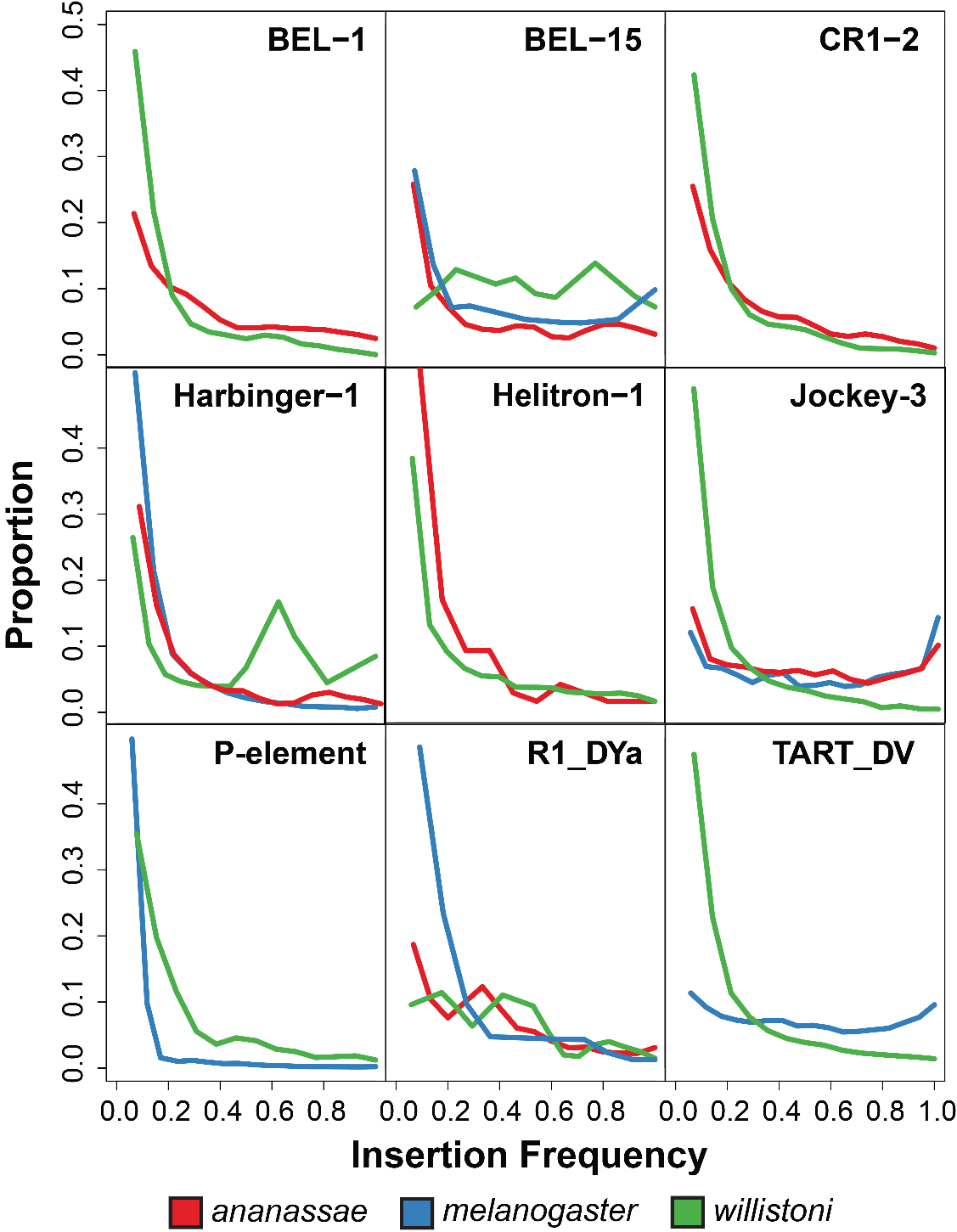
